# Computer-aided prediction of antigen presenting cell modulators for designing peptide-based vaccine adjuvants

**DOI:** 10.1101/232025

**Authors:** Gandharva Nagpal, Kumardeep Chaudhary, Piyush Agrawal, Gajendra P.S. Raghava

## Abstract

**Background:** Evidences in literature strongly advocate the potential of immunomodulatory peptides for use as vaccine adjuvants. All the mechanisms of vaccine adjuvants ensuing immunostimulatory effects directly or indirectly stimulate Antigen Presenting Cells (APCs). While numerous methods have been developed in the past for predicting B-cell and T-cell epitopes; no method is available for predicting the peptides that can modulate the APCs.

**Methods:** We named the peptides that can activate APCs as A-cell epitopes and developed methods for their prediction in this study. A dataset of experimentally validated A-cell epitopes was collected and compiled from various resources. To predict A-cell epitopes, we developed Support Vector Machine-based machine learning models using different sequence-based features.

**Results:** A hybrid model developed on a combination of sequence-based features (dipeptide composition and motif occurrence), achieved the highest accuracy of 96.91% with Matthews Correlation Coefficient (MCC) value of 0.94 on the training dataset. We also evaluated the hybrid models on an independent dataset and achieved a comparable accuracy of 94.93% with MCC 0.90.

**Conclusion:** The models developed in this study were implemented in a web-based platform VaxinPAD to predict and design immunomodulatory peptides or A-cell epitopes. This web server available at http://webs.iiitd.edu.in/raghava/vaxinpad/ and http://crdd.osdd.net/raghava/vaxinpad/ will facilitate researchers in designing peptide-based vaccine adjuvants.

## Introduction

Peptide subunit vaccines are hailed as an advancement over live or inactivated whole organism vaccines due to their ability to minimize adverse reaction [1]. Yet, antigenic peptides by themselves are poorly immunogenic since they lack the capability of activating the innate immunity. Activation of the innate immune system is required for stimulation of whole immune system including adaptive immunity. Hence, there is a need for inclusion of immunostimulants known as adjuvants in the subunit vaccine formulations. Conventionally, empirical approaches were used for adjuvant discovery, so far limited adjuvants have been approved and licensed for clinical use like alum, MF59, AS03 and AS04 [2].

Vaccine adjuvants effectuate their action by a variety of mechanisms with all of them involving the Antigen Presenting Cells (APCs) particularly the dendritic cells [3]. One of these mechanisms is the activation of the pattern recognition receptors (PRRs) on the APCs that recognize conserved microbial molecular signatures. PRR ligands shape the adaptive immune response mediated by the APCs. A majority of the vaccine adjuvants are ligands of PRRs making them potential targets for rational design of vaccine adjuvants [2]. Thus, hypothesis-driven adjuvant development relies on the expectation that the mechanistic understanding of the immune responses exhibited by PRR ligands would enable fine-tuning the specificity of adjuvants to attain vaccine efficacy and safety, simultaneously. An important example of a class of molecules that have been shown to have immunomodulatory effects and are poised to become safe and cost-effective adjuvants in future is – short immunomodulating peptides [4]. Figure 1 is a schematic representation of the adaptive immune cell activation by a coordination of antigen presentation to the naïve adaptive immune cell with the release of cytokine milieu mediated by PRR activation. Keeping in view the role of peptide ligands of PRRs in the activation of APCs, we introduce the term ‘A-cell epitopes’ for these immunomodulatory peptides.

**Figure 1:**
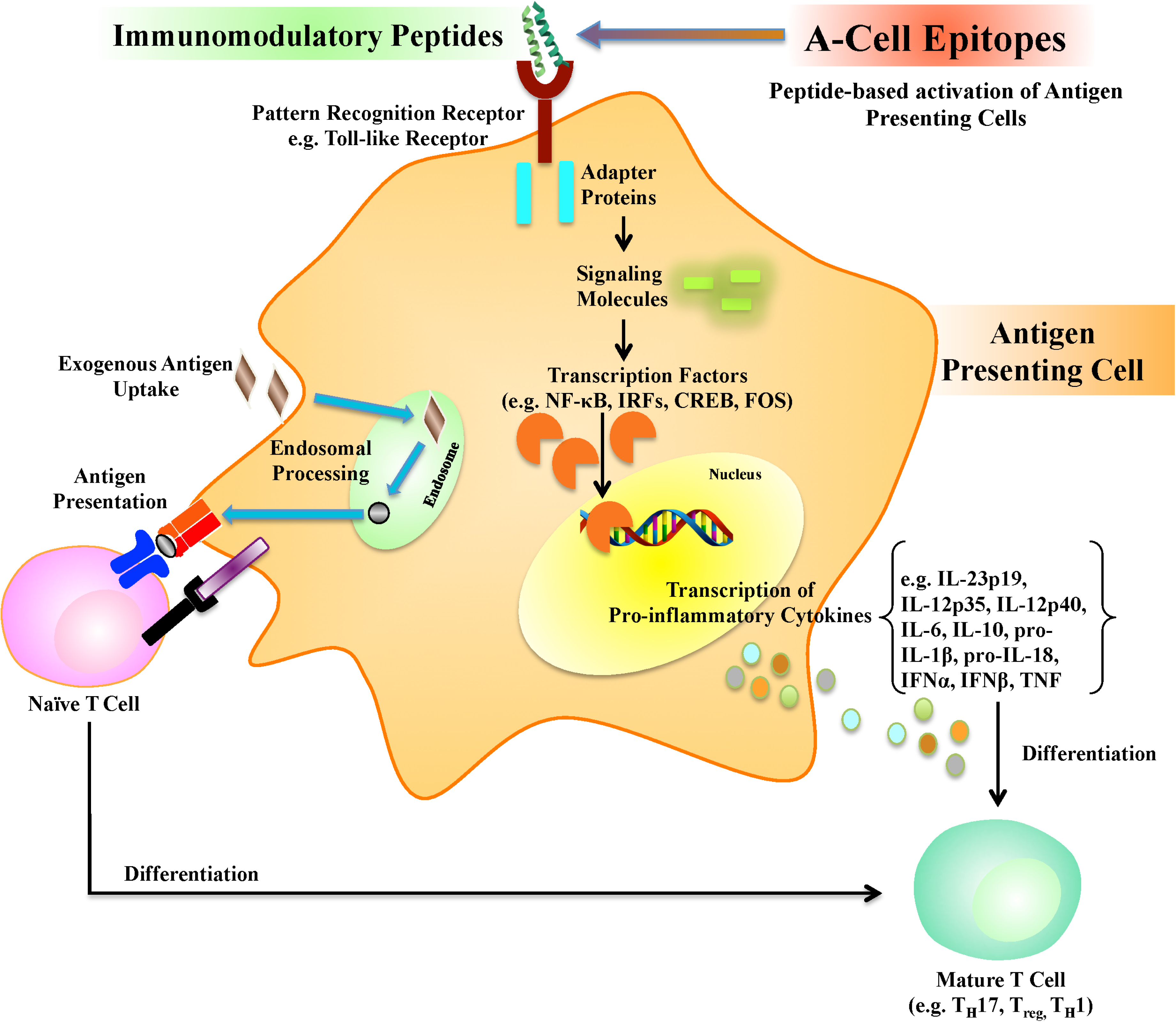
An illustrative mechanism of Antigen Presenting Cell (APC) activation caused by immunomodulatory peptides through innate immune receptors leading to the induction of adaptive immune cells. The immunomodulatory peptides are ligands of innate immune receptors that evoke cytokine expression through cellular signaling pathways. The cytokines lead to the maturation of naïve cells into mature adaptive immune cells such as various types of T-lymphocytes. Since the immunomodulatory peptides activate the APCs leading to the activation of the adaptive immune cells, they may be used as vaccine adjuvants and be called ‘A-cell epitopes’. The figure was drawn using ScienceSlides, made available at http://www.scienceslides.com/ by VisiScience.

Cationic Host Defense Peptides (HDPs) were originally discovered as antimicrobial peptides produced within the multicellular organisms having a broad-spectrum activity against bacteria, viruses, fungi, protozoa, etc [5]. Of late, HDPs and their synthetic analogs called Innate Defense Regulators (IDRs) have been realized to cause immunomodulatory effects like differentiation and activation of innate and adaptive immune cells, modulation of pro- and anti-inflammatory responses, chemo attraction, autophagy, apoptosis and enhancement of immune-mediated bacterial killing [6] **[25113635]**. Many Host Defense Peptides (HDPs) with known immunomodulatory effects are already in clinical trials [7]. IMMACCEL-R is a short synthetic peptide with immunomodulatory properties that has been commercialized for use as vaccine adjuvant in animals and birds for the purpose of antibody generation [8]. The human Cathelicidin antimicrobial peptide (CAMP or LL37) is another well-known antimicrobial peptide shown to induce immunomodulatory effects [9] and has been found to be associated with immune-related disorders like psoriasis [10] and morbus Kostmann [11].

Adjuvants have been incorporated into the vaccine formulations for qualitative alteration of the adaptive immune responses that are different from non-adjuvanted antigens. For instance, the adjuvants have been used to skew the immune responses with respect to Th1 (T-helper 1) cells versus Th2 cells, CD8+ versus CD4+ T cells, specific antibody types, etc. [2]. The PRR ligands produce these effects by virtue of the adapter proteins in the signaling pathways activated by the PRR. For example, bacterial flagellin protein causes adjuvant effect through TLR 5 and produces a mixed Th1 and Th2 response instead of polarized Th1 response and requires TLR adapter protein MyD88 for this effect. In contrast, monophosphoryl lipid A (MPL) and bacterial lipopolysaccharide (LPS) acting through TLR 4 activation lead to production of pro-inflammatory cytokine TNF leading to a polarized Th1 response instead of mixed Th1-Th2 response. While MPL signals through TRIF adaptor, LPS mediated activation of TLR 4 acts through both TRIF and MyD88 adapter proteins. Thus, MPL formulated on alum (AS04) stimulates a polarized Th1 cell response and is a component of licensed vaccine for HBV and papilloma that has proven to be both safe and effective.

Developing adjuvants based merely on empirical studies without the understanding of mechanisms is inadequate [2]. There is a need to develop systematic and rational approaches for designing highly potent vaccine adjuvants. One such approach could be the development of PRR ligands into vaccine adjuvants since their mechanism is known. In such a scenario, *in silico* models to screen and identify potential vaccine adjuvant candidates could prove to be useful as the existing experimental approaches are time and resources consuming [12]. Previously, our group developed a method, VaccineDA for predicting immunomodulatory oligodeoxynucleotides that can activate innate immune system via Toll-like receptor-9 (TLR-9). This tool can be used for designing oligodeoxynucleotide-based vaccine adjuvants as well as for genome-wide screening of vaccine adjuvants [13]. Recently, we also developed a method imRNA for designing single-stranded RNA (ssRNA) based vaccine adjuvants [14]. These methods may play an important role in designing DNA and RNA-based therapeutics as these methods allow a user to design oligonucleotides of desired immunogenicity.

In the last two decades, numerous method have been developed for predicting potential of peptides to stimulate adaptive arm of the immune system that include methods for predicting MHC binders, B-cell epitopes [15–23] and T-cell epitopes [24–31]. To the best of our knowledge, no method has been developed so far for predicting immunostimulatory potential of peptides to activate innate immunity. In this study, we made an effort to develop method for predicting immunomodulatory peptides that can activate innate arm of immune system or antigen presenting cells. These peptides activate the antigen presenting cells (e.g., dendritic, macrophages); hence, we propose that these immunomodulatory peptides be termed as ‘A-cell epitopes’.

In the present work, first we collected experimentally identified immunomodulatory peptides from the literature and included them in our positive set named A-cell epitopes. Next, we collected the human endogenously circulating peptides to build the negative set named A-cell non-epitopes. Combining the positive and the negative sets into a complete dataset, we developed Support Vector Machine (SVM) based computational models that can classify a new query peptide as A-cell epitope or non-epitope. To benefit the users of the scientific community, we provided the best performing SVM-based prediction models in the form of a web-based application called VaxinPAD to be used for identifying and designing novel A-cell epitopes. Such peptides identified computationally might serve as the starting molecules for designing peptide-based vaccine adjuvants.

## MATERIAL AND METHODS

### Dataset

The experimentally validated immunomodulatory peptide sequences were obtained from 16 patents. As an example of the sequences considered immunomodulatory, a set of sequences taken from a patent (US20110008318 A1), includes flagellin-derieved peptides that exhibit immunomodulatory effect by direct binding to TLR 5 as indicated by assays reporting increased NF-κB expression estimated from coupled luciferase activity and TNFα production estimated using flow cytometry. In one of the patents (US7462360 B2), a class of immunomodulatory peptides, called Alloferons, derieved from the bacteria challenged blood of larvae of the insect blowfly, *Calliphora vicina*., have been found to stimulate the cytotoxic anticancer activity of the human NK-cells and lymphocytes. In another case, a set of peptides as described in patent US8791061 B2 have been shown to enhance innate immunity by modulating the activity of type II transmembrane serine protease dipeptidyl peptidase (DPPIV) also known as CD26 or adenosine deaminase binding protein, expressed on major immune cells like activated T-Cells, B-Cells, NK-cells, macrophages and epithelial cells. With two major functions of signal transduction and proteolysis, the effects of DPPIV protein-mediated cellular processes include modulation of the chemokine activity by cleaving dipeptides from chemokine N-terminus that alters the receptor binding and specificity of the processed chemokine. DPPIV is a neutrophil chemorepellant and eosinophil chemoattractant too.

After removing the longer sequences, 304 unique sequences left in the length range of 3-30 residues were used to constitute the positive dataset named here as the A-cell epitopes. The upper bound of length 30 residues was kept as more than 90% of the originally collected epitope sequences were retained keeping this criterion used for removing very long sequences. In the absence of experimentally verified non-immunomodulatory peptides (non-epitopes), the experimentally identified endogenous human serum peptides [32, 33] were taken as non-epitopes. We assume these peptides are non-immunogenic as they are part of human serum, thus we assign them as non-epitopes. Only the sequences of the length 3-30 were taken into the negative dataset. In this manner, the Main dataset consisted of 304 A-cell epitopes and 385 non-epitopes. Table S1 (Additional File 1) provides the sequences and the source patent/publication for the positive and the negative datasets.

### Input Features

In order to develop any *in silico* model it is important to generate input features corresponding to each data point. In this study, a data point is the amino acid sequence of a peptide (either A-cell epitope or non-epitope). It is important to generate fixed length input features because machine-learning techniques require fixed length vector for developing a model. As the length of peptides is variable, thus we computed amino acid composition of A-cell epitopes and non-epitopes for developing models. We also computed the average amino acid composition of A-cell epitopes, non-epitopes and the human proteins, in order to understand compositional bias in A-cell epitopes. The amino acid composition for each sequence constituted the input vector of length 20, which was used for developing SVM-based prediction models. Similarly, the dipeptide composition vectors of length 400 were generated for A-cell epitopes and non-epitopes with each element of a vector corresponding to the composition value of each type of possible dipeptide. In addition to compositional features, we also generated binary features for developing models using fixed length of amino acids from terminus (N- or C- or both terminal) of peptides. In the case of binary feature, an amino acid is represented by a vector of 20, where the presence of amino acid is indicated by ‘1’ and the absence is presented by ‘0’ [26]. This means a peptide of length N is presented by a vector of length N*20 in the case of binary features.

### Motif Search

We used the Motif—EmeRging and with Classes—Identification (MERCI) Program[34][] to identify motifs exclusively occurring in the A-cell epitopes [35]. Though this program allows searching for gapped and ungapped motifs, but we restricted our analysis to the ungapped motifs. It is well established that in the case of T-cell epitopes, even a single residue mutation changes its immunogenicity [36] and can even eliminate the immunogenicity of the epitope [37]. Hence, intuitively the ungapped motifs found to be conserved among the positive sequences are more likely to help identify novel A-cell epitopes. Thus, we computed and compared the frequency of occurrence of ungapped motifs in A-cell epitopes, non-epitopes and the Swiss-Prot proteins.

### Support Vector Machine (SVM)

All the prediction models in this study were developed using SVM, which has been frequently used to develop models for epitope prediction in previous studies [15, 17, 27]. SVM has been the method of choice for building epitope prediction models especially T-cell epitopes [38] due to its ability to provide effective models on high dimensionality data with less data points. The dataset used in the current study contains data points comparable in number to the dimensionality. Hence, we optimized the prediction models on various parameters using the radial basis kernel of a freely available program SVM^*light*^ [39] to select the best performing models on different sets of features.

### Evaluation of models using internal and external validation

In this study, standard procedure was followed to evaluate the performance of models in order to avoid biases in performance due to over optimization. Our Main dataset was divided into two categories internal and external dataset, where the internal dataset contained ~80% sequences and the external dataset comprised of the remaining 20% sequences. In order to perform internal validation, we performed five-fold cross validation technique on internal dataset. In this technique, the dataset is divided in five sets, four sets are used for training a model, and the remaining set is used for testing the model. This process is repeated five times so each sequence is tested only one time. In order to perform the external validation of a model, the best model developed using five-fold cross validation is tested on an external dataset. It is important to assess the performance of a model on external or independent dataset because the performance of a model in internal validation may be biased due to optimization of the model [27]. The performance of models was measured using standard metrics namely Sensitivity, Specificity, Accuracy and Matthew’s Correlation Coefficient (MCC) [19, 35].

## RESULTS

### Compositional Analysis

One of the objectives of this study is to understand the nature of A-cell epitopes regarding the residues preferred in A-cell epitopes. Thus, we computed the average residue composition of A-cell epitopes and the non-epitopes. The non-epitope dataset consists of peptides occurring in the normal human serum assumed to be non-immunomodulatory. In addition, the average residue composition of the Swiss-Prot Human proteins was also computed and compared with that of the A-cell epitopes.In the A-cell epitope dataset, the percentage composition of an amino acid residue was calculated for each epitope, and the average of these values was plotted in Figure 2 for the corresponding amino acid. Similarly, the average percentage composition was calculated for all the amino acids in the non-epitope dataset and the Swiss-Prot Human proteins. As shown in Figure 2, the residues showing noticeable differences in average composition between A-cell epitopes and non-epitopes are C, D, E, I, L, R, S, T, V and W. Student’s t-test significance value (p-value) was calculated for each residue type to check whether the composition values among A-cell epitopes were different from those in the non-epitopes. In decreasing order of significance (increasing adjusted p-value), the residues R, E, T, S, D, V, W, L, I and C showed the most significant difference between the A-cell epitopes and non-epitopes among all of the residue types with adjusted p-values 1.76E-39, 1.66E-24, 7.04E-18, 3.59E-17, 3.72E-12, 2.79E-11, 5.21E-10, 7.91E-10, 1.24E-09, 1.76E-08 respectively (Additional File 1 Table S2). In particular, when compared to the human proteins taken from Swiss-Prot; R was found to have a higher average composition in A-cell epitopes. The average composition of R in non-epitopes is lower than that in the human proteins. Overall, the residues I, R, V and W were found to be more abundant in the A-Cell epitopes as compared to the non-epitopes and Swiss-Prot Human proteins.

**Figure 2:**
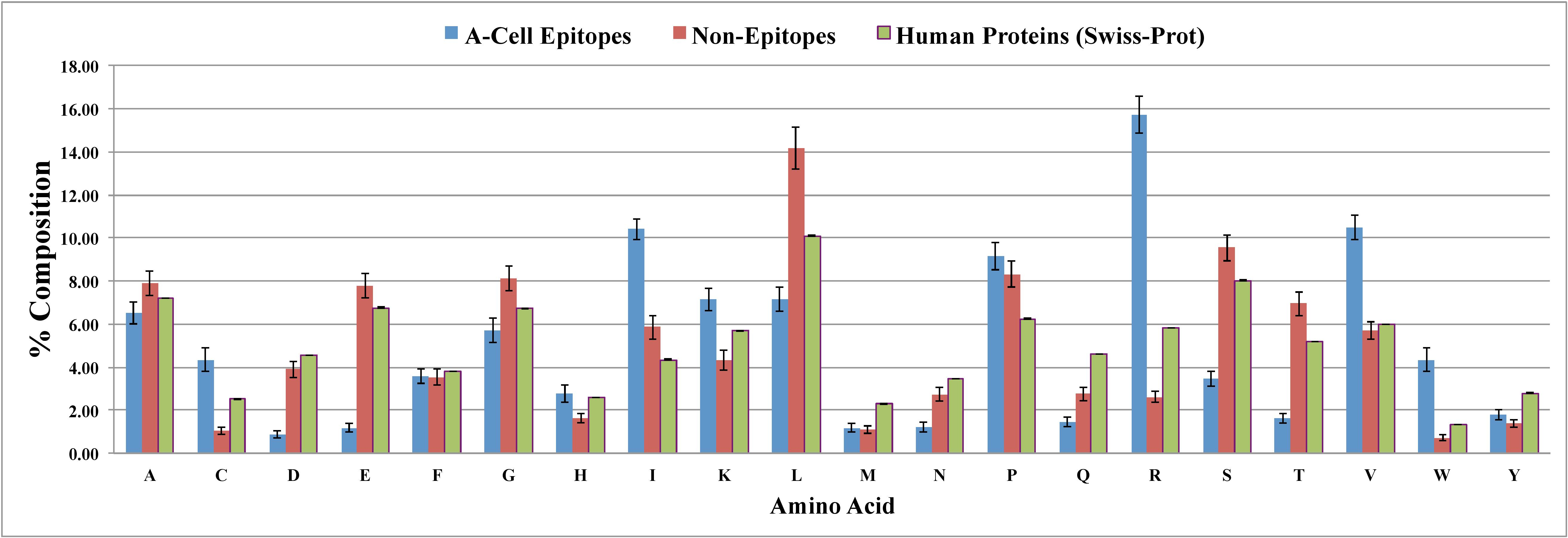
Bar plots showing the comparison of percent average amino acid composition of A-cell epitopes (blue) with non-epitopes (red) and Swiss-Prot Human proteins (green).

Similarly, the dipeptide and tripeptide compositions of the A-cell epitopes and non-epitopes were also compared with the Swiss-Prot Human proteins. Table S3 (Additional File 1) gives the average composition for each dipeptide in the A-cell epitopes, non-epitopes as well as the Swiss-Prot Human proteins. After sorting the table according to descending order of difference of dipeptide composition between the A-cell epitopes and the human proteins, top 10 dinucleotide include the residues I, R and V. But these motifs also contain other amino acids that show less significant difference of abundance as compared to the non-epitopes and human proteins. Similar analysis of tripeptide composition is shown in Table S4 (Additional File 1). In this case too, the top 10 tripeptide motifs include less abundant residues apart from I, R and V.

### Terminal Residue Preference

We performed position-specific analysis of residues in A-cell epitopes to understand the type of residues preferred at different positions in A-cell epitopes. In this study, Two-Sample Logo (TSL) tool (available at http://www.twosamplelogo.org/cgi-bin/tsl/tsl.cgi)] [40] was used to visualize residues preferred or not preferred in A-cell epitopes. Since the minimum peptide length in the dataset was 3, the N-terminal 3 residues of both the negative and the positive sequences were taken as input to build the N-terminus TSL. C-terminus TSL was obtained using the C-terminal 3 residues from the dataset. Figure 3 shows that the residues R, V and I are among the preferred residues in the A-cell epitopes at both the N and the C termini.

**Figure 3:**
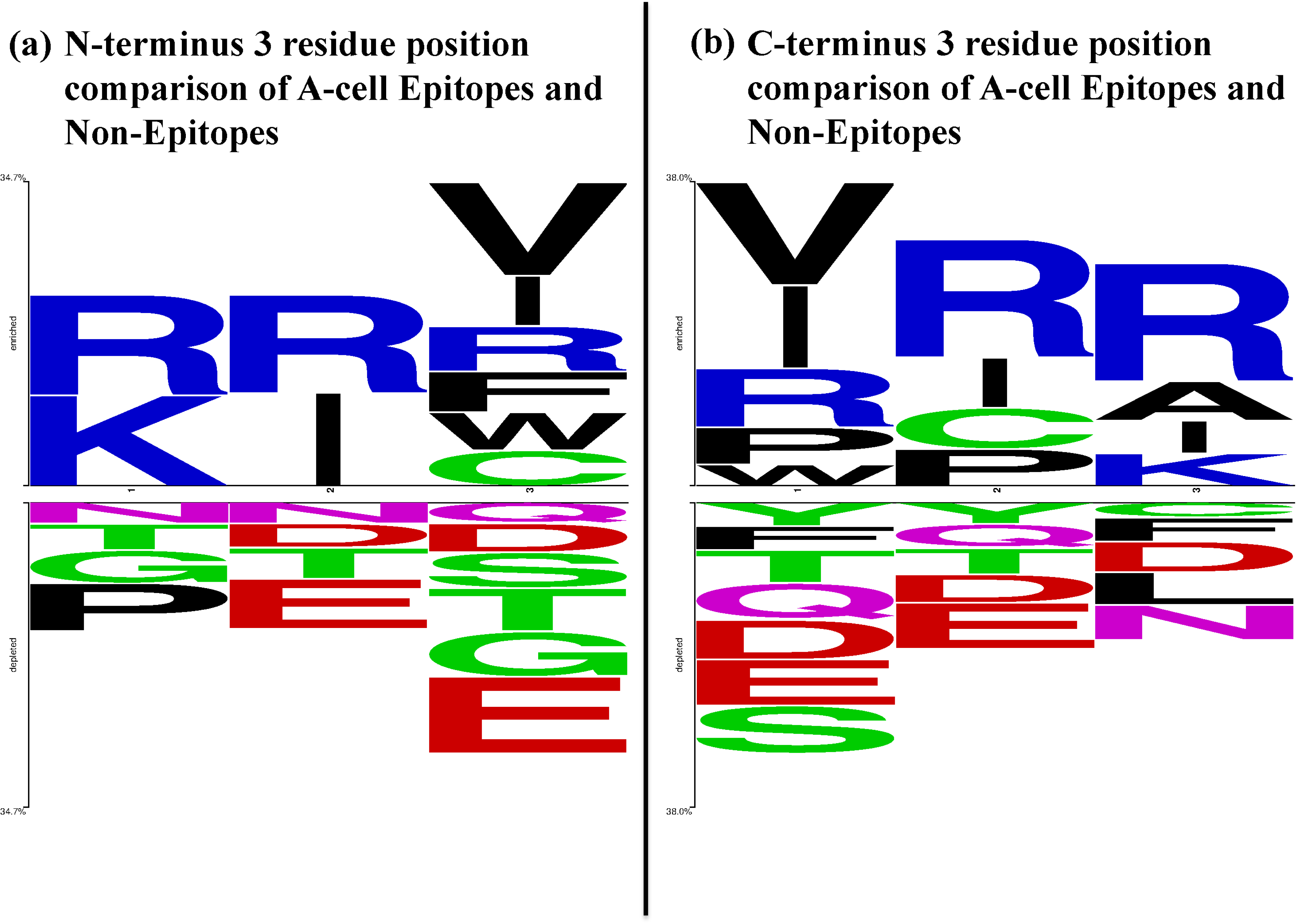
Two-Sample Logo of the 3 residue positions at the (a) N-terminus and (b) C-terminus of the A-cell epitopes and non-epitopes. Enriched label represents the positive dataset whereas depleted label represents the negative dataset. In a Two-Sample Logo, the height of a symbol at a residue position is proportional to the difference in symbol frequency between the positive and the negative datasets at that residue position. In the case of A-cell epitopes (as positives) and non-epitopes (as negatives), R is a preferred amino acid at terminal positions apart from I and V. The symbol colors are in accordance with the WebLogo default color scheme provided by the server available at http://www.twosamplelogo.org/cgi-bin/tsl/tsl.cgi. In the default WebLogo color scheme, residues G, S, T, Y and C appear in green color, N and Q are colored purple, K, R and H are depicted in blue, D and E are drawn red and P, A, W, F, L, I, M and V are shaded black.

### MERCI Motif Analysis

The Motif—EmeRging and with Classes—Identification (MERCI) Program is a software that helps in finding the motifs exclusive to one class when compared to another class of sequences. Table S5 (Additional File 1) provides the MERCI motifs exclusive to the A-cell epitopes as compared to non-epitopes. Top 10 ungapped motifs with respect to the occurrence in the A-cell epitope sequences have a frequent occurrence of I, R and V. On the other hand, ungapped MERCI motifs exclusive in non-epitopes (Additional File 1 Table S6) that are top 10 in abundance contain E, G, P and L.

### Rare Motif Occurrence

We compared the occurrence of peptide *n*-mers (*n*=3,4,5,6) in the A-cell epitopes and non-epitopes. First, the occurrence of each type of *n*-mer was counted in all of the Swiss-Prot proteins, and the *n*-mers were arranged in increasing order of occurrence. In this order, the *n*-mers were divided into 8 bins such that the 1st bin contained the *n*-mers least abundant in Swiss-Prot while the 8th bin contained the most abundant *n*-mers occurring in the Swiss-Prot. Next, the percentage of *n*-mers in a particular bin that occur in the Swiss-Prot was calculated with respect to the total number of *n*-mers in Swiss-Prot. Similar percentage value was calculated for A-cell epitopes and non-epitopes for each bin, and the values were presented in the form of a plot in Figure 4.

**Figure 4:**
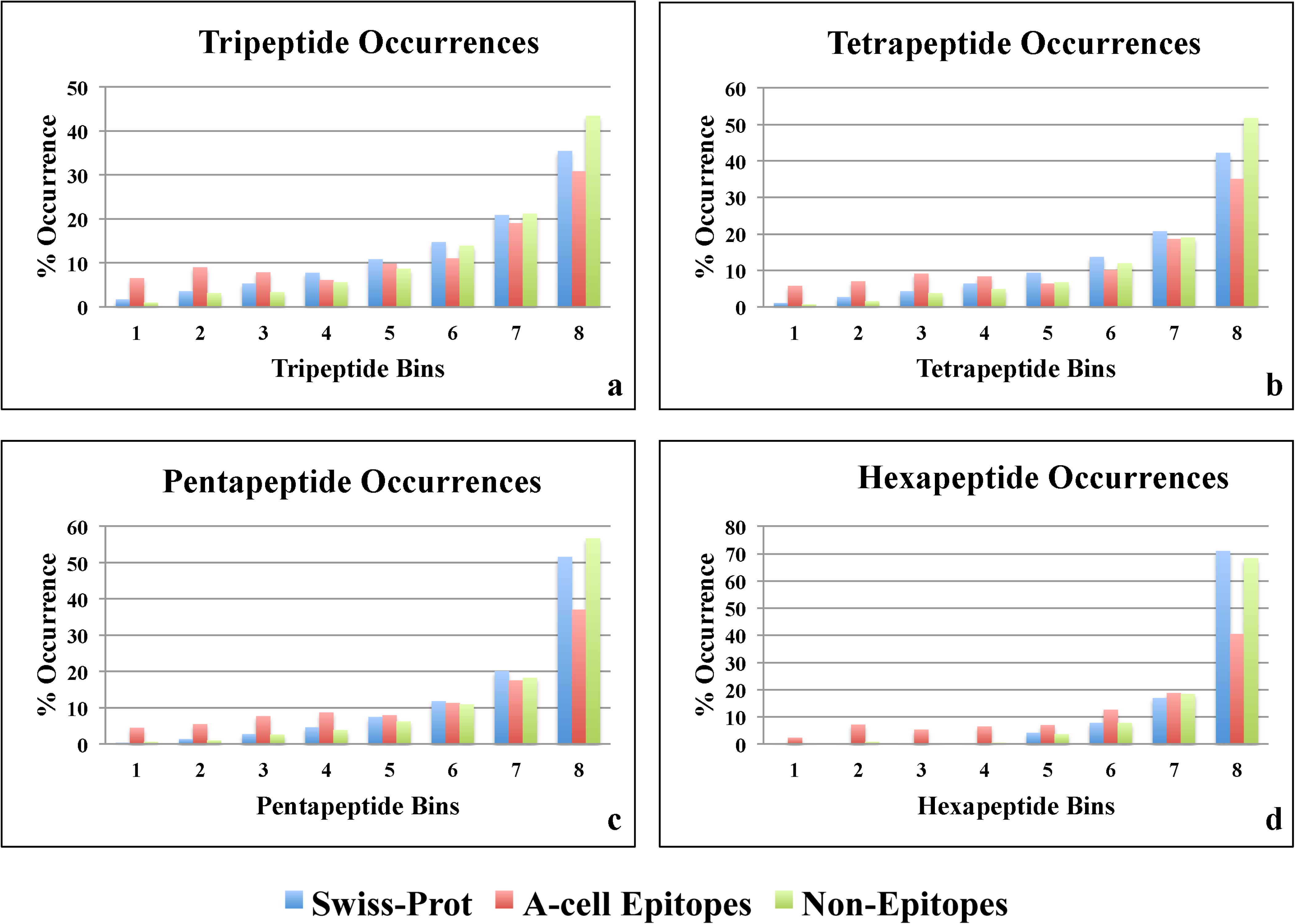
Comparison of occurrence of (a) tripeptides, (b) tetrapeptides, (c) pentapeptides and (d) hexapeptides divided into 8 bins in the ascending order of occurrence (most rarely occurring to most abundant) in Swiss-Prot proteins.

Figure 4(a) shows that tripeptides in the first three bins occur more in A-cell epitopes while those of 4^th^, 5^th^, 6^th^, 7^th^ and 8^th^ bin (most abundant Swiss-Prot tripeptides) occur more in the non-epitopes. For tetrapeptides (Figure 4(b)), the bins having more number of tetrapeptides occurring in the A-cell epitopes than non-epitopes are 1^st^, 2^nd^, 3^rd^ and 4^th^. Figure 4(c) shows the occurrence of the pentapeptides. The bins having distinctly more pentapeptides in the A-cell epitopes than non-epitopes are again the first four bins. On the other hand, the percentage occurrence of hexapeptides of A-cell epitopes is lower than non-epitopes and Swiss-Prot proteins only in the 8^th^ bin (Figure 4(d)).

### Prediction of Immunomodulatory Peptides

The sequence-based analyses like residue composition preferences; position-wise residue preference and motif search indicated that these features could help in discriminating the A-cell epitopes from non-epitopes. We developed SVM-based prediction models using SVM^*light*^ by from the dataset of 304 A-cell epitopes as positive sequences and 385 non-epitopes as negative sequences. From each of the positive and negative datasets, ~80% sequences were kept in the Training-Testing dataset while the remaining ~20% were kept in the Independent Dataset. Thus, the Training-Testing dataset had 243 positive and 308 negative sequences. The prediction models were developed in three categories.

#### Composition-based Models

The amino acid composition (AAC) and dipeptide (DPC) composition were used to develop SVM-based models on the Training-testing Dataset.On evaluating the performance parameters, the AAC model gave an accuracy of 95.10% and the Matthews Correlation Coefficient (MCC) value of 0.90 as given in Table 1. The DPC model also gave a similar performance in terms of accuracy (95.64%) and MCC (0.91) values. When the AAC and DPC models on the terminal 5 residues of the sequences individually (N or C terminus – Nter, Cter) or together (N and C termini combined - NCter) were developed, the NCter AAC and DPC models performed closest to but not better than the AAC and DPC models with MCC values for NCter AAC being 0.88 and NCter DPC 0.87 (Table 1).

**Table 1:** The performance of SVM-based models developed using various features including different types of amino acid composition. This is internal cross-validation where models were evaluated on training dataset using five-fold cross-validation.

| **Features** | **Threshold** | **Sensitivity(%)** | **Specificity(%)** | **Accuracy(%)** | **MCC** |
| --- | --- | --- | --- | --- | --- |
| AAC | 0.00 | 94.24 | 95.78 | 95.10 | 0.90 |
| Nter AAC | 0.00 | 89.08 | 91.50 | 90.34 | 0.81 |
| Cter AAC | 0.00 | 91.27 | 92.31 | 91.81 | 0.84 |
| NCter AAC | -0.10 | 92.58 | 95.14 | 93.91 | 0.88 |
| DPC | 0.00 | 95.06 | 96.10 | 95.64 | 0.91 |
| Nter DPC | -0.10 | 86.03 | 86.64 | 86.34 | 0.73 |
| Cter DPC | -0.10 | 89.96 | 94.33 | 92.23 | 0.84 |
| NCter DPC | 0.10 | 92.14 | 94.33 | 93.28 | 0.87 |
| N5 bin | -0.10 | 87.34 | 88.26 | 87.82 | 0.76 |
| C5 bin | -0.20 | 86.46 | 89.47 | 88.03 | 0.76 |
| N5C5 bin | 0.00 | 89.96 | 91.90 | 90.97 | 0.82 |
| N10 bin | -0.20 | 86.37 | 85.88 | 86.21 | 0.72 |
| C10 bin | -0.20 | 83.19 | 88.14 | 86.21 | 0.71 |
| N10C10 bin | -0.30 | 88.50 | 90.40 | 89.66 | 0.78 |
| AAC+motif | 0.00 | 96.30 | 96.43 | 96.37 | 0.93 |
| DPC+motif | 0.00 | 95.47 | 98.05 | 96.91 | 0.94 |

#### Binary models

Binary models take the residue position into account by representing each residue type as a binary vector. We considered 5 and 10 residue positions from either end of the peptide sequences (N and C termini) and developed SVM-based models individually and in combination. In case of 5-residue position consideration, the model developed on combined 5 residue positions on both the N and C termini (N5C5 bin in Table 1) performed the best giving an accuracy value 90.97% and MCC value 0.82 while the N10C10 bin model performed the best among the 10-residue position models (accuracy 89.66% and MCC 0.78).

#### Hybrid Model

In the previous motif analysis, motifs exclusive to the A-cell epitopes were identified using the MERCI program. We checked whether the motif information added to composition could help improve the performance of prediction. Indeed the AAC+motif model that combined the information of presence of motifs exclusive to A-cell epitopes with amino acid composition achieved a better performance than AAC model on the Training-Testing dataset giving an accuracy value of 96.37% and MCC value of 0.93 (Table 1). Yet, the best model among all the feature combinations was that of motif information combined to the DPC (DPC+motif model) that gave an accuracy of 96.91% and the MCC value 0.94.

### Performance of the models on the independent datasets

The independent dataset consisted of ~20% of the total dataset resulting in 61 positive sequences and 77 negative sequences. The performances of the AAC and DPC models on the independent dataset were comparable to those on the Training-Testing dataset with AAC model giving an accuracy of 92.75% and MCC of 0.85 while DPC model giving an accuracy of 95.65% and MCC of 0.91 (Table 2). The MCC values of the NCter AAC and NCter DPC models in the independent dataset evaluation (0.92 and 0.90 respectively) were also close to those found in the Training-Testing Dataset evaluation (0.88 and 0.87 respectively). Similar to the Training-Testing Dataset results, the N5C5 bin and N10C10 bin models performed the best (MCC 0.93 and 0.81 respectively) among the binary models on the independent dataset. The hybrid model (AAC+motif and DPC+motif) gave accuracies 90.58% and 94.93% respectively, while the MCC values found were 0.81 and 0. 90 respectively when evaluated on the independent dataset (Table 2). Figure 5 is a plot of the MCC values of various SVM models on the Training-Testing Dataset shown as bar plot along with the MCC values of the models obtained on the Independent Dataset drawn as the line graph. The MCC values on both the datasets are comparable for each of the models developed indicating that the models are not over-optimized on the Training-Testing Dataset.

**Figure 5:**
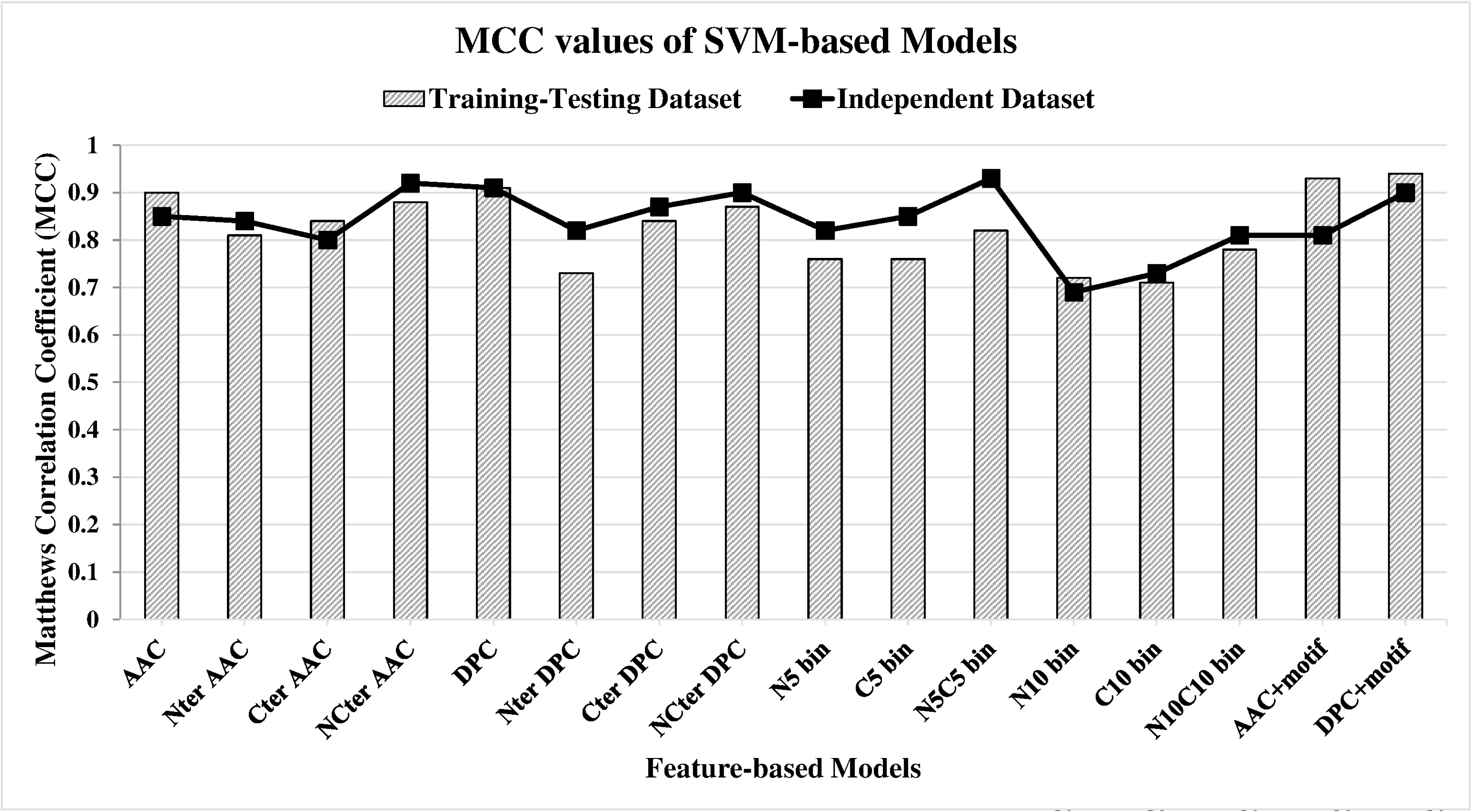
Comparison of the Support Vector Machine-based prediction models on the Training-Testing and the Independent Datasets. The striped bars correspond to the Matthews Correlation Coefficient (MCC) values obtained for the models on the Training-Testing Dataset and the solid line joins the MCC values of the models on the Independent Dataset. For each model, the MCC values for the Training-Testing Dataset and the Independent Dataset are comparable indicating the reliable prediction capabilities of the models.

**Table 2:** The performance of A-cell epitope prediction models developed using different features on an independent dataset (external cross-validation).

| Features | Threshold | Sensitivity(%) | Specificity(%) | Accuracy(%) | MCC |
| --- | --- | --- | --- | --- | --- |
| AAC | 0.00 | 91.80 | 93.51 | 92.75 | 0.85 |
| Nter AAC | 0.00 | 85.45 | 96.92 | 91.67 | 0.84 |
| Cter AAC | 0.00 | 89.09 | 90.77 | 90.00 | 0.80 |
| NCter AAC | -0.10 | 92.73 | 98.46 | 95.83 | 0.92 |
| DPC | 0.00 | 95.08 | 96.10 | 95.65 | 0.91 |
| Nter DPC | -0.10 | 90.16 | 92.21 | 91.30 | 0.82 |
| Cter DPC | -0.10 | 96.36 | 90.77 | 93.33 | 0.87 |
| NCter DPC | 0.10 | 95.08 | 94.81 | 94.93 | 0.90 |
| N5 bin | -0.10 | 87.27 | 93.85 | 90.83 | 0.82 |
| C5 bin | -0.20 | 92.73 | 92.31 | 92.50 | 0.85 |
| N5C5 bin | 0.00 | 96.36 | 96.92 | 96.67 | 0.93 |
| N10 bin | -0.10 | 92.31 | 79.59 | 84.00 | 0.69 |
| C10 bin | -0.20 | 80.77 | 91.84 | 88.00 | 0.73 |
| N10C10 bin | -0.30 | 96.15 | 87.76 | 90.67 | 0.81 |
| AAC+motif | 0.00 | 90.16 | 90.91 | 90.58 | 0.81 |
| DPC+motif | 0.00 | 93.44 | 96.10 | 94.93 | 0.90 |

\* **MCC**: Matthews Correlation Coefficient, **AAC**: Amino Acid Composition, **DPC**: Dipeptide Composition, **Nter AAC**: N terminus (5 residues) amino acid composition, **Cter AAC**: C terminus (5 residues) amino acid composition, **NCter AAC**: N and C terminus (5 residues each) amino acid composition, **Nter DPC**: N terminus (5 residues) dipeptide composition, **Cter DPC**: C terminus (5 residues) dipeptide composition, **NCter DPC**: N and C terminus (5 residues each) dipeptide composition, **N5 bin**: Binary profile 5 residues from N-terminus, **C5 bin**: Binary profile 5 residues from C-terminus, **N5C5 bin**: Binary Profile 5 residues from N-terminus and C terminus, **N10 bin**: Binary profile 10 residues from N-terminus, **C10 bin**: Binary profile 10 residues from C-terminus, **N10C10 bin**: Binary Profile 10 residues from N-terminus and C terminus, **AAC+motif**: amino acid composition with MERCI motif score, **DPC+motif**: dipeptide composition with MERCI motif score.

### VaxinPAD: The web Interface for prediction of A-cell epitopes

For providing the SVM-based A-cell epitope prediction methods developed in the present study to the scientific community, we designed an *in silico* platform VaxinPAD (available at http://crdd.osdd.net/raghava/vaxinpad). The platform has a couple of utilities that may help the user in designing peptide-based adjuvants as well enhance or diminish the immunomodulatory potential of the query sequence.

Designing of vaccine adjuvants: The ‘PREDICTION’ module of the VaxinPAD platform allows the user to check whether the query sequence would be immunomodulatory on the basis of SVM score. It allows the virtual screening of the A-cell epitopes among a library of input peptide sequences.

#### Immunomodulatory regions in a protein

The ‘PROTEIN ADJUVANTS’ module of VaxinPAD does a window search across the length of a query protein sequence to identify immunomodulatory patches, the window size being user-defined. LL37, a well-known immunomodulatory peptide is a 37 amino acid peptide derived from human Cathelicidin. This menu may help the researchers in identifying more such peptides that are immunomodulatory.

#### Peptide Sequences Dataset

Finally, VaxinPAD includes a menu ‘DATASET’ that is a list of immunomodulatory peptides collected from literature. Among the sequences in the database only the peptides of length 3-30 were used for development of prediction models in the current work.

## DISCUSSION

Previously, peptide-based vaccine adjuvants were largely being developed as ligands of innate immunity receptors like TLR-4 and TLR-2 [41] or as self-assembling nanostructure forming entity [42]. Recently, it has been realized that short immunomodulatory peptides can be developed as potential vaccine adjuvants [4].Cationic Host Defense Peptides were previously known to have antibacterial activity by direct killing of the pathogen [43]. Of late, these peptides have been found to evoke the innate immunity by a variety of mechanisms [6]. A majority of these mechanisms involve pattern recognition receptors (PRRs) playing important roles especially in the Antigen Presenting Cells (APCs) like Dendritic cells, macrophages, etc. Since these peptides activate APCs, we call these peptides as ‘A-cell epitopes’ (Antigen Presenting Cell epitopes). The A-cell epitopes undertaken in the present study were collected from the patent literature that included host defense peptides as immunomodulatory sequences. To the best of the authors’ knowledge, the present study is the first attempt to develop an *in silico* tool for designing immunomodulatory peptides as the first step towards engineering novel peptide-based vaccine adjuvants.

An important finding in this study was that the residues preferred in A-cell epitopes include arginine (R). Arginine enrichment of the peptide sequences is an important aspect of increasing the cellular uptake of cell-penetrating peptides (CPPs) [44]. Cathelicidins are recognized as an important class of host defense peptides that includes many arginine-rich peptides [45]. Further, human Cathelicidin-derived peptide LL37 that is rich in basic residues arginine and lysine has been reviewed as a promising immunomodulatory peptide with cell penetrating properties [46]. Hence, sequence analysis of the A-cell epitopes may indicate cell-penetrating ability to be an associated property of the A-cell epitopes.

Another aspect of our sequence analysis is the occurrence of *n*-mers found sparsely in the naturally occurring proteins. Patel *et al*, 2012 [47] found that peptide pentapeptides occurring rarely in the universal proteome when introduced into the end of the antigenic sequence enhanced its antigenicity and also suggested that on exogenous addition these rare pentapeptides could act as immunomodulators and thus could be developed as adjuvants. In our analysis too, we found that tripeptides, tetrapeptides and pentapeptides occurring rarely in Swiss-Prot proteins to be present more in the A-cell epitopes than non-epitopes. This fits well with the intuition that the immune system is more likely to react to rarely encountered sequence motifs than frequent ones.

On evaluating the performance of SVM models based on composition, the dipeptide composition showed no improvement over amino acid composition. The binary models also showed a lower performance than composition-based models. On the other hand, addition of the motif information increased the performance of both the amino acid composition model as well as dipeptide composition to achieve the maximum accuracies of ~96%.

The peptides designed using the tools developed in the present study might act by various mechanisms and receptors for activating the innate immunity owing to the fact that the training dataset of the prediction models contains peptides acting by diverse cell signaling routes. Hence, the *in silico* tool presented here could help an investigator to begin with a choice of peptides that may be the starting points in the development of vaccine adjuvants. Whether these peptides actually prove to be useful as adjuvants, would have to be tested experimentally. Another limiting aspect of the present study is the exclusion of very long immunomodulatory sequences. With larger datasets and receptor-specific ligands made available in future, studies subsequent to the present investigation might help design peptides eliciting a specific desired immune response. Nonetheless, VaxinPAD developed in this study for predicting the immunomodulatory peptides sets a stage for the advancement of rational peptide-based vaccine adjuvant designing.

*In silico* methods for predicting and identifying DNA and RNA-based immunomodulatory molecules have already been developed [13, 14]. The current study is the first attempt to develop models for predicting immunomodulatory peptides for the development of vaccine adjuvants. Though other biomolecules like lipopolysaccharides and glycosaminoglycans also cause activation of innate immunity by binding to the PRRs, the literature currently does not hold a sufficient number of molecules for developing prediction models in these categories. Future studies might focus on the development of *in silico* tools for predicting such immunomodulatory biomolecules for obtaining new vaccine adjuvants. In addition to this, peptides with non-natural chemical modifications might offer better adjuvants too. Correspondingly, computational tools for prediction of modified peptides might also become an area of development.

## CONCLUSION

Host Defense Peptides have been realized as promising immunomodulators likely to become potential vaccine adjuvants [43]. With immunomodulatory properties, novel peptides predicted to be A-cell epitopes using the models developed here are also likely to have potential to provide host protection against pathogens. Many Host Defense Peptides (HDPs) with known immunomodulatory effects are already in clinical trials [43]. Despite the associated toxicity of the A-cell epitopes due to their pleiotropic effects on the immune system, rational design of Innate Defense Regulators (IDRs) that are synthetic analogs of HDPs is in pressing demand for having immunopotentiators with reduced toxicity and increased specificity of immune responses. We have developed SVM-based models for prediction of A-cell epitopes that could be used to formulate vaccine adjuvants. These models have been implemented in the form of webserver VaxinPAD available at http://webs.iiitd.edu.in/raghava/vaxinpad/ and http://crdd.osdd.net/raghava/vaxinpad/ freely to the scientific community.

## ABBREVIATIONS

APC: Antigen Presenting Cell
PRR: Pattern Recognition Receptor
HDP: Host Defense Peptide
IDR: Innate Defense Regulator
TLR: Toll-like receptor
CAMP: Cathelicidin Antimicrobial Peptide
MHC: Major Histocompatibility Complex
SVM: Support Vector Machine
MERCI: Motif—EmeRging and with Classes—Identification
TSL: Two-Sample Logo
CPP: Cell Penetrating Peptides

## AUTHOR CONTRIBUTION

PA and GN collected the data. PA, KC and GN organized the data. KC, GN and PA performed the experiments. GN and KC developed the web interface. GN, KC, PA and GPSR analyzed the data. GN and GPSR prepared the manuscript. GPSR conceived the idea and coordinated the project.

## ACKNOWLEDGEMENTS

Authors are thankful to the projects, Open Source Drug Discovery (OSDD) and GENESIS BSC0121. GN and KC are thankful to CSIR and PA is thankful to the Department of Science and Technology (DST), Government of India for providing fellowships.

## Availability of data and materials

The datasets used for this study are provided at http://webs.iiitd.edu.in/raghava/vaxinpad/sequences.php and http://crdd.osdd.net/raghava/vaxinpad/sequences.php as well as Additional File 1 Table S1.

## Ethics approval and consent to participate

Not required in this study.

## Consent for Publication

Not required

## Additional File

### Additional File 1

**Table S1**: Dataset (A-cell epitopes and Non-epitope) used for training, testing and validation; including suorce of information. **Table S1**: Dataset (A-cell epitopes and Non-epitope) used for training, testing and validation; including suorce of information. **Table S3**: The percent dipeptide composition and difference in composition in A-cell epitopes, non-epitopes and human proteins. **Table S4**: The percent tripeptide compositions of A-cell Epitopes, Non-epitopes and Human proteins. Rows are sorted in decreasing order of values in the 5th column. **Table S5**: Motifs exclusively found in A-cell epitopes and their frequency of occurrence, sorted in the decreasing order of counts. **Table S6**: Motifs exclusively found in Non-epitopes and their frequency of occurance, sorted in decreasing order of counts.

